# Age-related decrements in cortical gyrification: Evidence from an accelerated longitudinal dataset

**DOI:** 10.1101/2020.03.05.979344

**Authors:** Christopher R. Madan

## Abstract

Cortical gyrification has been found to decrease due to aging, but thus far this has only been examined in cross-sectional samples. Interestingly, the topography of these age-related differences in gyrification follow a distinct gradient along the cortex relative to age effects on cortical thickness, likely suggesting a different underlying neurobiological mechanism. Here I examined several aspects of gyrification in an accelerated longitudinal dataset of 280 healthy adults aged 45-92 with an interval between first and last MRI session of up to 10 years (total of 815 MRI sessions). Results suggest that age changes in sulcal morphology underlie these changes in gyrification.

## Introduction

Over the past decade, brain imaging studies have demonstrated several gradients in activation related to functional networks of regions (e.g., Margulies et al., 2016; Murphy et al., 2019; Sormaz et al., 2018). Distinct cortical gradients in structural properties of the brain also exist, such as in cortical thickness (e.g., Hogstrom et al., 2013; Madan & Kensinger, 2018; Salat et al., 2004; Sowell et al., 2003) and myelination (e.g., Carradus et al., 2020; Glasser & Van Essen, 2011; Grydekand et al., 2013; Mangeat et al., 2015; Shafee et al., 2015). However, one of the most identifiable characteristics of the human brain is its folding structure. Despite macro-scale consistencies between individuals, everyone’s cortical folding pattern is unique. While these folding patterns change due to aging, the underlying process of this change is not well understood.

Elias and Schwartz (1969) developed a procedure to quantify cortical gyrification as a ratio between the area of the total cortical surface relative to the ‘exposed’ surface. Here a stereological approach was used, though was limited by available technologies. This methodology was later refined to use the pial and outer contour lengths estimated from coronal sections by Zilles et al. (1988). The outlines shown in Figure 1A display this visually, with the pial length shown in blue and the outer contour length shown in green. The ratio of these lengths provide a measure of cortical folding, referred to as the gyrification index. A brain with a lower gyrification index will be smoother and less folded; a rodent’s brain has a much lower gyrification index than a human brain. Extending this procedure to 3D surfaces, Schaer et al. (2008, 2012) developed an automated approach for calculating cortical gyrification from a surface mesh, through the generation of an outer surface that encloses the gray matter surface generated by FreeSurfer (as shown in Figure 1B) along with the corresponding surface measure of gyrification (Figure 1C).

**Figure 1.**
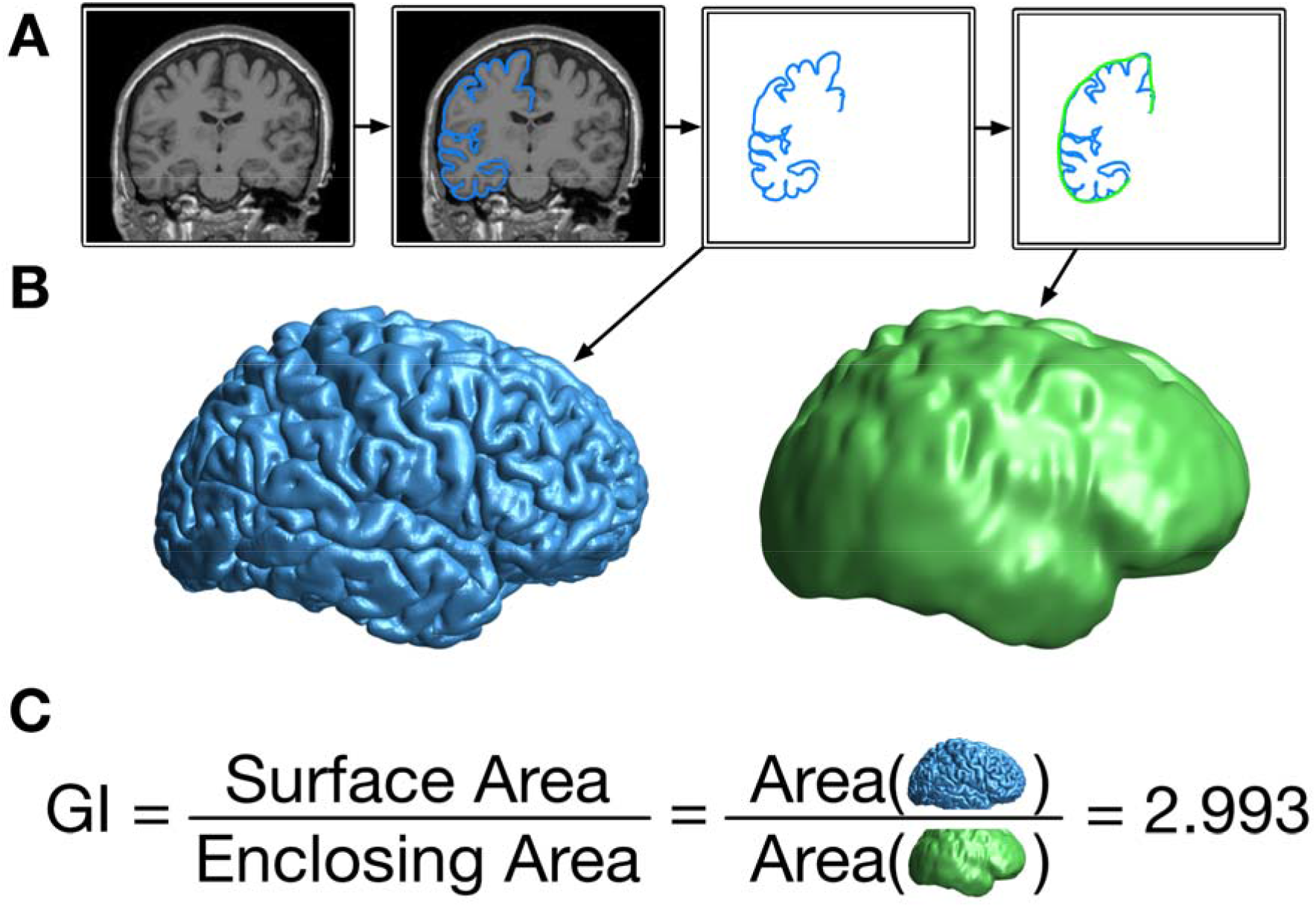
Illustration of the calculation procedure for global gyrification. (A) From the original T1-weighted volume, the outer contour of the gray matter is traced, and then surrounded by a smooth enclosing surface. (B) When done as a 3D surface, this results in the pial (blue) and enclosing (green) surfaces. (C) The gyrification index (GI) is the ratio of the cortical surface area divided by the surface area of the enclosing surface.

As initially shown by Zilles et al. (1988) and replicated in more recent studies (e.g., Hogstrom et al., 2013; Madan & Kensinger, 2016), gyrification is relatively highest in the parietal and temporal lobes. Several studies have shown that global cortical gyrification decreases with age in several cross-sectional samples (Cao et al., 2017; Hogstrom et al., 2013; Lamballais et al., 2020; Madan & Kensinger, 2016, 2018; Madan, 2018). It has also been shown that the topography of these changes is distinct from cortical thickness, where gyrification primarily decreases in the parietal lobe (e.g., Hogstrom et al., 2013; Madan & Kensinger, 2016, 2018). Cortical thickness, in contrast, primarily decreases in frontal and temporal regions—with age related decreases associated with changes in cortical myelination affecting gray/white matter contrast as well as decreases in the dendritic density of pyramidal neurons (Dickstein et al., 2007; Duan et al., 2003; Hao et al., 2007; Peters, 2002), mechanisms distinct from age changes in gyrification. Further demonstrating the utility of gyrification measurements, studies of patient populations have demonstrated differences in gyrification relative to healthy individuals in relation to Alzheimer’s disease (King et al., 2010; Liu et al., 2012), schizophrenia (Cao et al., 2017; Palaniyappan & Liddle, 2012; Palaniyappan et al., 2015), autism (Schaer et al., 2013), and major depression disorder (Cao et al., 2017), among other psychiatric disorders. Within healthy samples, age-related differences in cortical folding structure have also been associated with individual functional differences (e.g., see Lamballais et al., 2020; McDonough & Madan, in press).

Using cross-sectional data, global gyrification has previously been estimated to decrease by approximately 0.035 GI/decade (Madan & Kensinger, 2016; Madan, 2018), though there is also a consideration of measurement validity. For example, in an analysis of test-retest reliability, where ten sessions were conducted just 2-3 days apart (each), a mean within-participant deviation of 0.04 was found (Madan & Kensinger, 2017). Moreover, plots of cross-sectional data reveal a large amount of off-axis variability (i.e., age-unrelated) indicating that, though reproducible, the link between gyrification and aging is relatively weak and influenced by many other factors. Based on these considerations, a longitudinal dataset with a larger interval between scans would be prudent for evaluating the influence of aging on cortical gyrification.

Here I examine changes in cortical gyrification in an accelerated longitudinal sample to directly examine cortical folding changes with age at the individual level, rather than in cross-sectional datasets. While accelerated longitudinal samples have previously been used to examine age changes in other morphological measures, such as cortical thickness (e.g., Storsve et al., 2014), this has yet to be done with gyrification. It is currently unclear how gyrification changes with aging—for instance, is the decrease in gyrification in the parietal lobe more evident because the parietal lobe has the highest gyrification? Moreover, does the distribution of gyrification change with age?

## Methods

### Dataset

Data from 280 healthy adults (aged 45-92) from the OASIS-3 (Open Access Series of Imaging Studies 3) dataset (LaMontagne et al., 2019) were included in the analyses presented here. From the full dataset, participants were only included in the present analyses if four conditions were met. (1) There must be at least two MRI sessions available with T1-weighted volumes collected. (2) MRI data had to be collected using either the Siemens TIM Trio 3 T scanner or Siemens BioGraph mMR PET-MR 3 T scanner (less than 5% of the otherwise available data were collected using one of three other MRI scanners). (3) There had to be an interval from first to last MRI session spanning at least three years. (4) From the clinical sessions, there had a Clinical Dementia Rating (CDR) score of zero at every assessment. All raw data is available from https://www.oasis-brains.org.

Across the 280 included adults, the Mini-Mental State Exam (MMSE) was administered in a total of 1904 clinical sessions, with between 2 and 20 clinical sessions per participant (*M±SD* = 6.70*±*3.21 administrations). Of the 1904 administrations, only four yielded an MMSE score below 25. For two individuals, these sub-25 MMSE scores were later followed by higher MMSE scores (here, scoring at least 29) on multiple subsequent administrations. For the remaining two individuals, these were the last MMSE scores recorded (for these specific individuals, there were 5 and 9 MMSE administrations, respectively).

The interval between the first and last MRI session acquired for each participant ranged from 3 to 10 years (*M±SD* = 5.47±1.91 years). Of the 280 participants, 117 had two MRI sessions, 96 had three sessions, 45 had four sessions, and 22 had five or more sessions (to a maximum of seven). In total, 1138 T1-weighted volumes from 815 MRI sessions were examined. If more than one T1-weighted volume was acquired during an MRI session, all volumes were processed (see procedure below) and the gyrification estimates for the scans were averaged. The same protocol was used for the other derived measures.

Structural data was collected using two different Siemens 3 T MRI scanners with MPRAGE sequences: (1) 581 of the 815 MRI sessions were collected with a Siemens TIM Trio 3 T scanner, with parameters: TR=2.4 ms; TE=3.2 ms; flip angle=8°; voxel size=1×1×1 mm. (2) The remaining 234 MRI sessions were collected with a Siemens BioGraph mMR PET-MR 3 T scanner, with parameters: TR=2.3 ms; TE=3.0 ms; flip angle=9°; voxel size=1×1×1 mm.

### MRI Processing

All T1-weighted volumes were processed using FreeSurfer v6.0 on a machine running RedHat 4.8.5-16 (Fischl, 2012; Fischl & Dale, 2000; Fischl et al., 2002). (Note, these are not the same FreeSurfer outputs as publicly distributed from OASIS, which were estimated using older versions of FreeSurfer.) All T1-weighted volumes were processed independently with the standard FreeSurfer pipeline (i.e., `recon-all`), i.e., not using the longitudinal processing pipeline, to allow the individual scan derived measures to be comparable in reliability to previous cross-sectional studies.

Gyrification index was calculated using FreeSurfer, as described in Schaer et al. (2008, 2012) and illustrated in Figure 1. This process involves generating an enclosing surface which serves the same purpose as the outer contour used by Zilles et al. (1988) but exists in 3D space. This enclosing surface involves a parameter of how ‘tight’ to bridge across gyri, as opposed to falling into sulci. The default settings were used, based on validation work conducted by Schaer and colleagues.

Sulcal morphology, width and depth, was estimated for 16 major sulci, using the calcSulc toolbox (Madan, 2019a). The sulci are the central, post-central, superior frontal, inferior frontal, parieto-occipital, occipito-temporal, middle occipital and lunate, and marginal part of the cingulate sulci, in both the left and right hemispheres. Analyses here are only conducted on the mean sulcal width and depth, averaging across the 16 sulci.

#### Anterior-posterior gradient in gyrification

The standard FreeSurfer pipeline results in variation in the number of surface mesh vertices for each participant. To adjust for this, gyrification measurements for each MRI session were resampled to the common surface space using FreeSurfer’s spherical surface co-registration (mris_preproc; Fischl et al., 1999). This registration allows for vertex-wise comparisons in gyrification across participants, as shown in Figure 2A.

**Figure 2.**
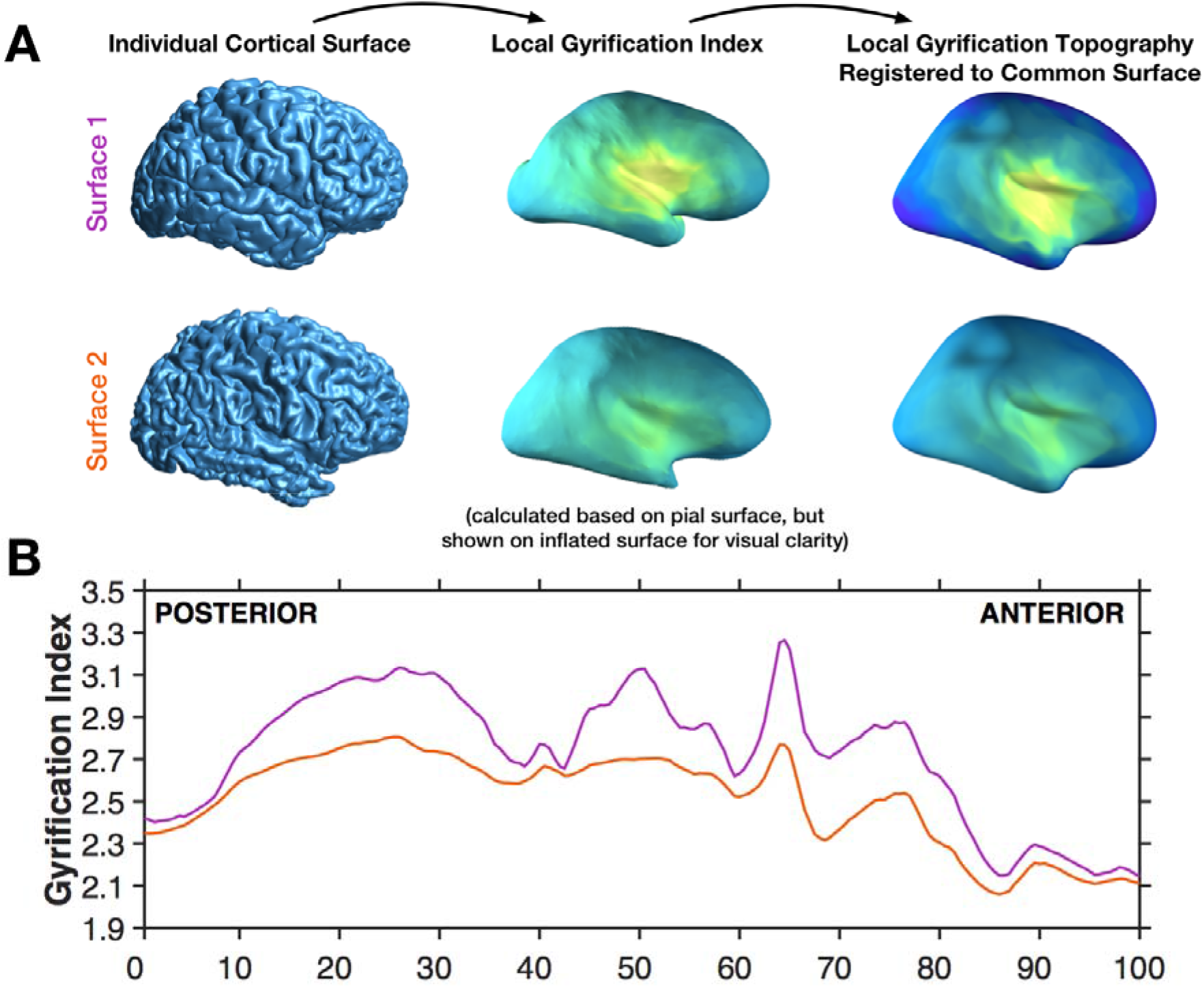
Illustration of the calculation of the anterior-posterior gradient. (A) Individual brain pial surfaces are used to generate local gyrification index topography (based on Schaer et al., 2012), these are then resampled to the common space, through registration of the individual pial surface to the FreeSurfer standard surface. (B) The local gyrification index is then averaged across vertices for 200 coronal sections, shown for the same two example surfaces as in panel A.

This common surface space was then divided into 200 coronal sections, analogous to the contour procedure used by Zilles et al. (1988). This allows for mean gyrification to be simplified into an anterior-posterior plot, as demonstrated in Figure 2B. Note that, however, vertex-wise gyrification measures were first estimated based on the Schaer et al. (2012) procedure that relies on 25-mm radius local region of interest. This results in a gyrification index value for each vertex of the surface mesh, referred to as a local gyrification index. Gyrification for each section is spatially autocorrelated with adjacent sections. To-date there do not appear to be any publications that have examined the anterior-posterior gyrification pattern using the FreeSurfer gyrification calculation implementation (i.e., the Schaer et al. [2012] approach).

### Statistics

Age-related changes in global gyrification were examined as the slope of decline in gyrification with age. In subsequent analyses, the relative contribution of sulcal width and depth, in explaining the age decrements in gyrification are evaluated using a mediation analysis.

## Results

### Global gyrification

Before examining the topography of gyrification, I evaluated age changes in global gyrification, which itself is yet to be examined in a longitudinal dataset. The decline in gyrification was estimated using a linear mixed effects model to estimate the slope (allowing for random intercepts for each participant; i.e., different starting points, but a common slope). As shown in Figure 3, the fixed effect of age was statistically significant [*t*(813)=14.41, *p*<.001] with a decreasing slope of 0.04291 GI/decade [95% C.I.= 0.03499—0.04603].

**Figure 3.**
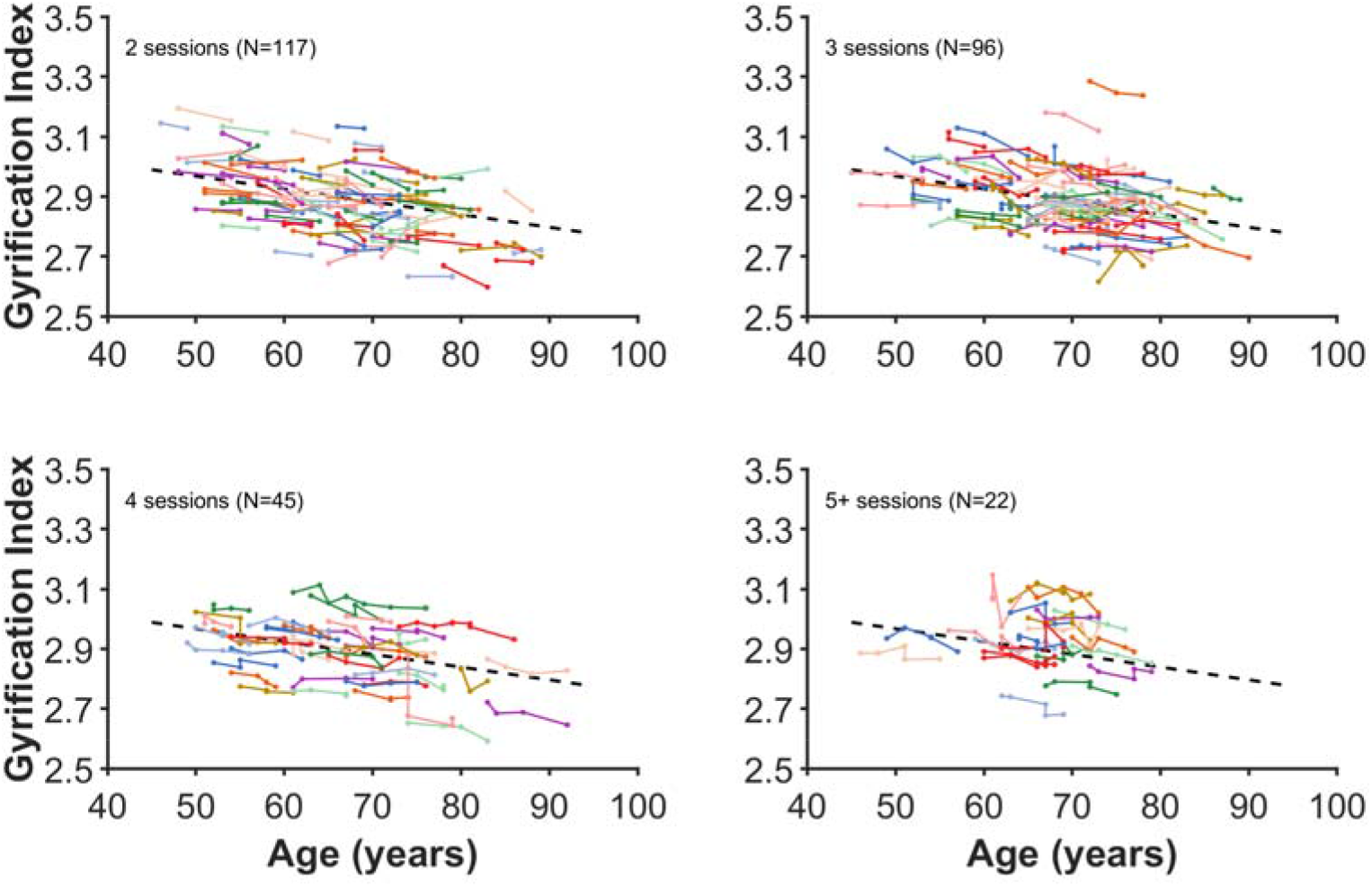
Age changes in global gyrification. Each coloured line represents an individual participant, with each dot corresponding to an MRI session. The plot has been divided into panels for each set of timepoints to improve readability. Dashed line present in all panels show the decrease in global gyrification based on all data, estimated using the linear mixed effects model.

For comparison, a similar analysis was conducted on fractal dimensionality, another measures of cortical structure. The results of this analysis are reported in the Appendix.

### Anterior-posterior gradient in gyrification

The results of this analysis are shown in Figure 4. Looking from anterior to posterior, the gradient gradually rises to a peak mid-way through the frontal lobe (approx. percentile 75), followed by a relative plateau through the section that subtends the temporal lobe, with a trough as the anterior-posterior section is increasingly represented by the parietal lobe (percentile 37). A higher plateau peak subtends the parietal lobe, and then gradually declines as gradient transitions into the occipital lobe (beginning from around percentile 15).

**Figure 4.**
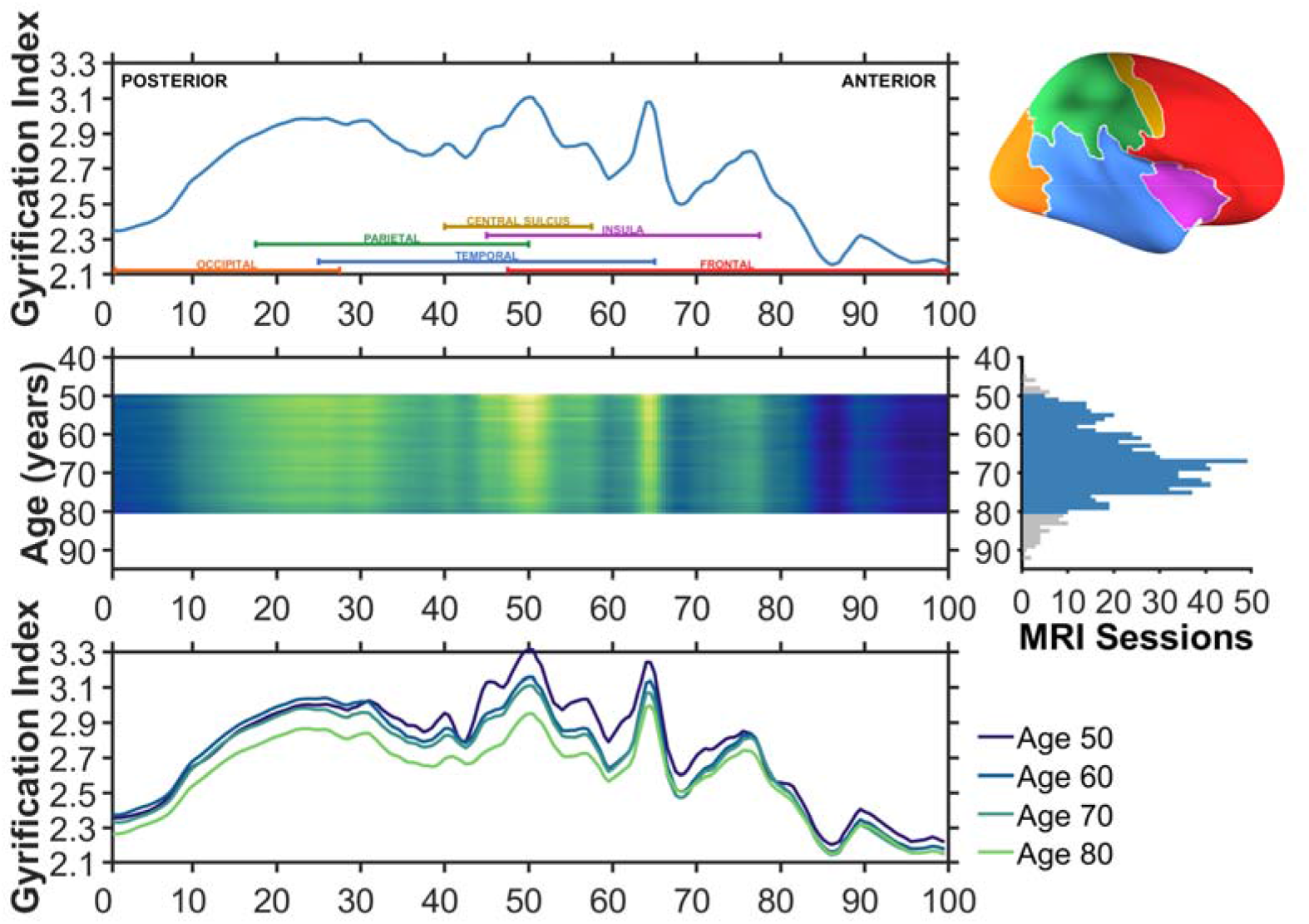
Gyrification index across posterior (percentile 0; caudal) to anterior (percentile 100; rostral). The upper row illustrates the average gyrification across the common surface space. Coloured labels denoting each of the four lobes (frontal: red; parietal: green; temporal: blue; occipital: orange), insula (purple), and central sulcus (yellow) are included to aid in interpretation. An inflated cortical surface is shown on the right to help visualise the relative extents of these regions, though a folded brain was used for the actual analyses. The middle row shows the mean anterior-posterior gyrification for each age between 50 and 80, inclusive; brighter colours denote regions of higher gyrification index. This subset of ages was selected based on having sufficient sessions per age to use in the estimation, as shown in the right panel. A total of 746 of the available 815 MRI sessions were in this age range. The lower row shows the mean anterior-posterior gyrification for individuals aged 50, 60, 70, and 80. Aging is associated with overall decreases in gyrification, but these appear to be most pronounced in the parietal lobe.

The middle and lower rows of Figure 4 show that while gyrification decreases globally, they are most pronounced in the parietal lobe. Further examination of the surface topography can better differentiate gyrification in parietal and temporal regions. These aging results demonstrate that the overall distribution of gyrification does not change with age, it merely diminishes in magnitude.

### Topography of gyrification

As gyrification is calculated as the ratio of areas of the cortical surface to an enclosing surface, the gyrification index is highest at the insula—as shown Figure 5A. Here I found a similar pattern in the present accelerated longitudinal dataset, as shown in Figure 5B. The decline in gyrification is most pronounced in the parietal lobe and posterior aspects of the frontal lobe. However, on individual cortical surfaces, even examining changes over nearly a decade, changes in the surfaces are only barely visible (see Figure 6).

**Figure 5.**
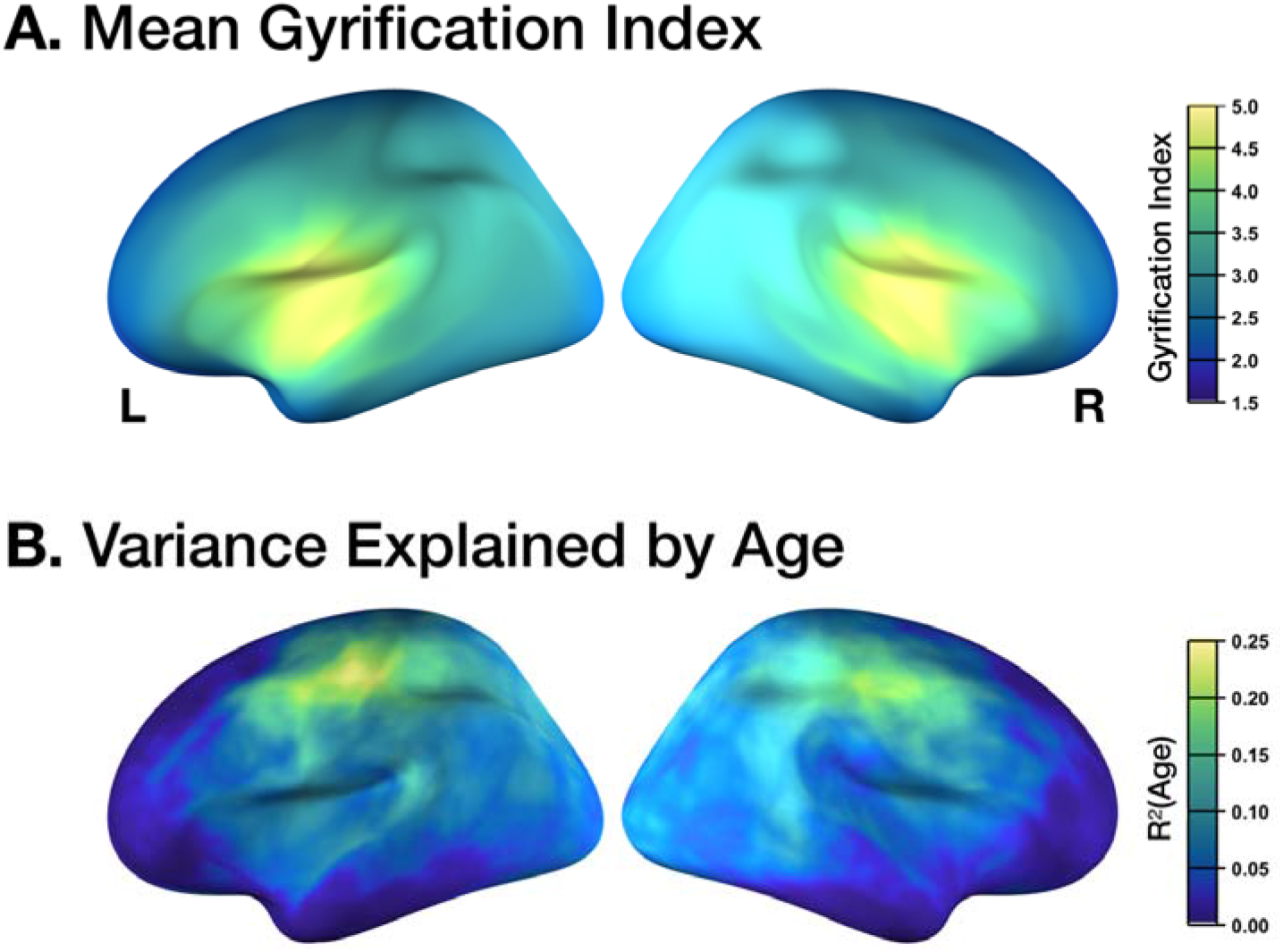
Topography of gyrification. (A) Mean gyrification, with the highest values corresponding to the insula. (B) Age changes in gyrification, with the highest values corresponding to the parietal lobe and posterior aspects of the frontal lobe (i.e., primary motor and somatosensory regions). Values determined based on vertex-wise regression of local gyrification index with age.

**Figure 6.**
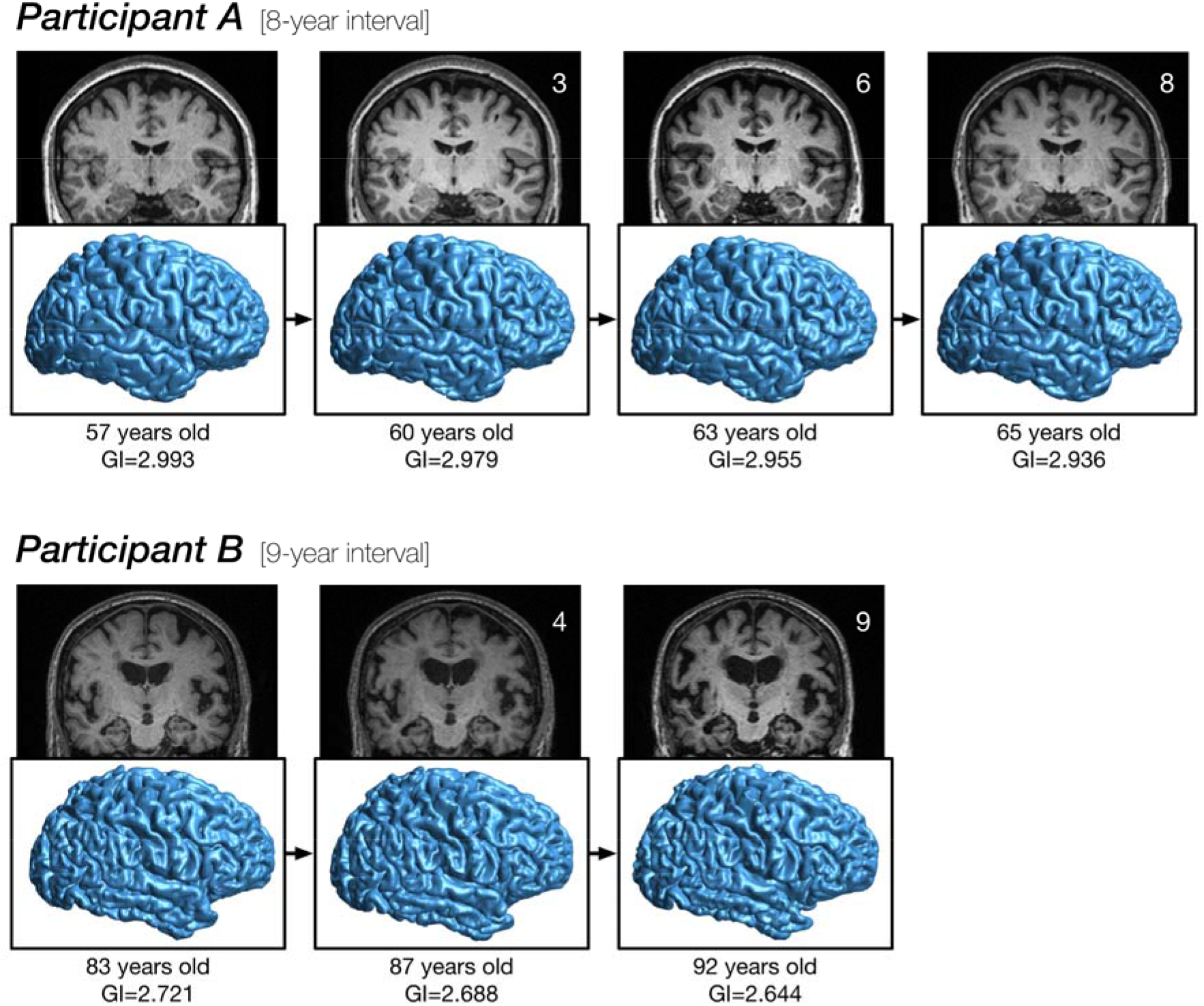
Longitudinal age changes in cortical folding. Cortical surface reconstructions for two participants, illustrating changes in folding over nearly a decade each. GI denotes gyrification index. See Madan (2015) for details related to rendering the cortical surfaces.

### Sulcal prominence

Figure 6 shows that the overall pattern of cortical folding remains relatively consistent, despite small reductions in the gyrification index. Visible from the MRIs themselves, however, does indicate changes in the sulcal width and depth. Here I examined how well global measures of sulcal morphology, both width and depth, can alternatively explain variability in gyrification. To test this, I conducted a multi-level mediation analysis on the longitudinal global gyrification index measurements, with random intercepts for each participant and based on all available timepoints.

Standardised parameter estimates for the model are shown in Figure 7, along with all associated standard error (SE) estimates in parentheses. Age was significantly related to increases in sulcal width [*a_1_* = 0.463 (0.028); *t*(813)=16.41, *p*<.001] as well as decreases in sulcal depth [*a_2_* = −0.362 (0.022); *t*(813)=16.76, *p*<.001]. Gyrification index was significantly related to both mediators, i.e., sulcal width and depth, and a significant direct effect of age remained after the mediators were modelled [sulcal width: *b_1_* = −0.105 (0.029); *t*(813)=3.69, *p*<.001; sulcal depth: *b_2_* = 0.676 (0.027); *t*(813)=25.12, *p*<.001; remaining direct effect of age: *c’* = −0.071 (0.020); *t*(813)=3.52, *p*<.001].

**Figure 7.**
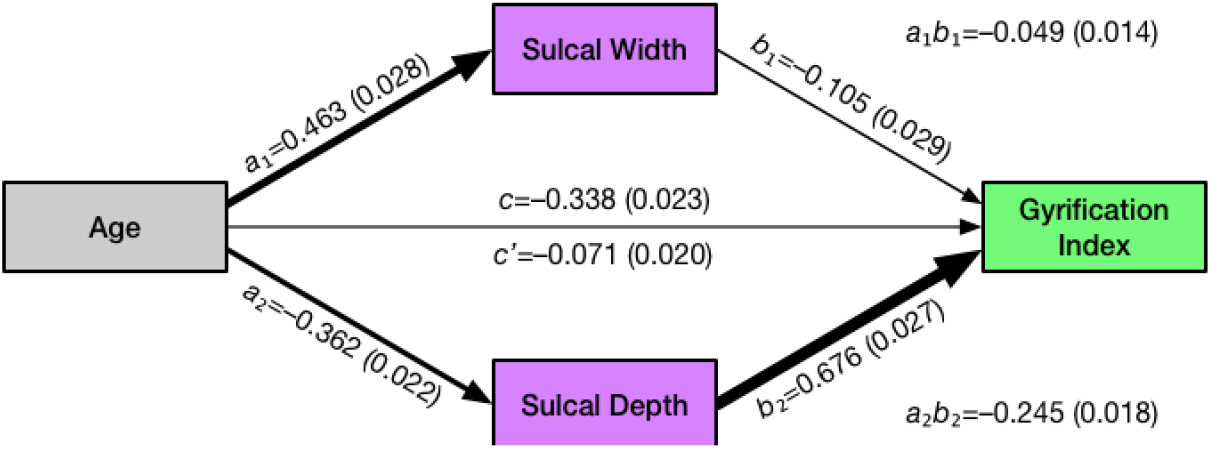
Path diagram for age effects on gyrification index, considering the mediators of sulcal width and depth. Standardised parameter estimates for the model are shown for each effect, along with standard error (SE) estimates in parentheses.

Mediation analyses indicated that both sulcal width and depth mediated the effects of age on gyrification index [sulcal width: *a_1_b_1_* = −0.049 (0.014); Sobel’s *Z*=3.59, *p*<.001; sulcal depth: *a_2_b_2_*= −0.245 (0.018); Sobel’s *Z*=13.94, *p*<.001]. The entire mediation model accounted for 49.9% of the variability in gyrification (i.e., *R^2^*). Proportion-mediated effect size estimates demonstrated that most of this variability was accounted for by sulcal depth (36.2%), with lesser proportions being explained by sulcal width (7.2%) and the remaining direct effect of age (10.6%). Note that these summed values slightly exceed the total amount of variability explained, as sulcal width and depth are not orthogonal. These results indicate that age related decreases in sulcal depth largely account for the apparent changes in gyrification, and both sulcal width and depth are more sensitive to effects of aging than the gyrification index measure.

## Discussion

From the global gyrification and anterior-posterior analyses, it is clear that the age decreases in gyrification are gradual. Moreover, the anterior-posterior analysis was designed to evaluate if these age changes in gyrification were related to shifts in the underlying distribution of cortical folding, but this does not appear to be the case. Instead, there is a global decrease, with some regions more affected than others, as highlighted in the topography analyses. Converging with prior cross-sectional studies, regions most affected by age related gyrification changes are distinct from those regions affected by changes in cortical thickness.

While studies examining cross-sectional data suggest that global gyrification involves a large degree of age-independent variability, the results of the longitudinal analyses here show a relatively consistent age decline, distinct from differences in the y-intercept of the gyrification index (i.e., Figure 3). Though changes in gyrification due to aging have not previously been investigated using an anterior-posterior gradient approach, the overall gradient (i.e., the upper row of Figure 4) is relatively consistent with Zilles et al. (1988, Fig. 6). The overall topography, as shown in Figure 5, appears consistent with previously published results (e.g., Schaer et al., 2008; Cao et al., 2017; Lamballais et al., 2020). As discussed in the Introduction section, previous studies have demonstrated age differences in gyrification, but as of yet, this has only been in cross-sectional samples (Cao et al., 2017; Hogstrom et al., 2013; Lamballais et al., 2020; Madan & Kensinger, 2016, 2018; Madan, 2018).

Despite numerous prior studies reporting decreases in gyrification with age, these studies provide little towards explaining the underlying mechanism. While a mechanism is not presented here either, the current results provide some insight into a more specific characterisation of how aging influences cortical folding. Many prior studies of brain morphology have indicated that cortical surface area is not affected by aging (e.g., Hogstrom et al., 2013; McKay et al., 2014; Storsve et al., 2014). This lack of influence of age on cortical surface area measurements also rules out any systematic changes in the minor deformations in the folds along the gyri. More specifically, these minor folds could be measured using other approaches, such as indices of the spatial frequency/power spectra of cortical folding as providing information distinct from gyrification itself (as in Madan, 2019b). Age-related changes in cortical thickness (and volume) are often found, but follow from a distinct topography than have been reported for gyrification (e.g., Hogstrom et al., 2013; Madan & Kensinger, 2016, 2018; McKay et al., 2014). However, long before neuroimagers began to examine cortical thickness as an index of cortical atrophy, radiologists have qualitatively assessed sulcal prominence—alternatively referred to as widening, enlargement, or dilation—as a key measure of age-related atrophy (Coffey et al., 1992; Drayer, 1988; Huckman et al., 1975; Jacoby et al., 1980; Laffey et al., 1984; LeMay, 1984; Pasquier et al., 1996; Roberts & Caird, 1976; Scheltens et al., 1997; Tomlinson et al., 1968; Turkheimer et al., 1984; Yue et al., 1997). Indeed, by visual inspection, cortical atrophy is much more readily assessed from sulcal features than from cortical thickness (see Figure 6).

Using current automated methods, sulcal morphology can also be quantitatively measured, where the width and depth of major sulci can be identified and estimated (e.g., Kochunov et al., 2005; Madan, 2019a). Across a number of studies and samples (albeit generally with much smaller samples), sulcal morphology has been reliably associated with age-related differences (Jin et al., 2018; Kochunov et al., 2005; Li et al., 2011; Liu et al., 2010, 2013; Madan, 2019a; Rettmann et al., 2006; Shen et al., 2018). The work presented here provides additional specificity in how gyrification appears to decrease with age, a step towards understanding the underlying mechanism. Here I show that gyrification changes can be considered strongly related to changes in the underlying sulcal morphology, particularly depth. However, despite this well-established relationship between sulcal morphology and aging, the neurobiological mechanism underlying this change in the fundamental organisation of cortical folding remains unclear and a topic for further study.

More generally, this work adds to the growing literature demonstrating that the availability of open-access MRI data can facilitate advances in our understanding of brain morphology well beyond the goals of the researchers that originally collected the data (see Madan, 2017, for an overview of benefits and considerations). It is worth acknowledging that the results presented here likely underestimate the extent that gyrification decreases with age. Older adults that are interested and able to participate in a research study in their 80s and 90s are very likely to be a biased sample of individuals for their age cohort, demonstrating better physical and mental health than many of their peers (i.e., an issue of external validity; see Pearl & Bareinboim, 2014, and Keyes & Westreich, 2019, for more nuanced discussions).

## Abbreviations

CDR: Clinical Dementia Rating
FD: fractal dimensionality
fMRI: functional magnetic resonance imaging
GI: gyrification index
MMSE: Mini-Mental State Exam
MRI: magnetic resonance imaging
OASIS: Open Access Series of Imaging Studies

## Data accessibility

All raw data is available from https://www.oasis-brains.org.

## Acknowledgments

I would like to thank Jonathan Reardon for feedback on an earlier version of this manuscript. This work would not have been possible without data from OASIS-3 and the funding that supported it (Principal Investigators: T. Benzinger, D. Marcus, J. Morris; NIH P50AG00561, P30NS09857781, P01AG026276, P01AG003991, R01AG043434, UL1TR000448, R01EB009352). I would also like to acknowledge the support of NVIDIA Corporation with the donation of a Titan Xp GPU that was used in this research.

## Appendix: Longitudinal changes in global cortical fractal dimensionality

Madan and Kensinger (2016) demonstrated that fractal dimensionality can be a more sensitive measure of age-related differences in cortical structure than conventional measures, including gyrification. To provide a complementary analysis, although tangential to the focus of the current study, here I present the age-related longitudinal changes in fractal dimensionality, from the same data as in the other analyses. To-date, longitudinal analyses of fractal dimensionality have not yet been conducted, and these results serve as an initial comparison between the gyrification results (e.g., participants in each panel are in the same line colour as in Figure 3). As expected from prior work, Figure A1 shows that there is less off-axis variability (i.e., random y-intercepts) with fractal dimensionality than are consistently found with gyrification. As with gyrification, a linear mixed effects model with random slopes for each participant was calculated, with a significant age fixed effect [*t*(813)=25.55, *p*<.001] with a decreasing slope of 0.01325 FD/decade [95% C.I. = 0.01223—0.01426].

As fractal dimensionality is a summary statistic of the complexity of a structure, a topographical analysis cannot be equivalently carried out as for gyrification—though fractal dimensionality can be calculated for parcellated cortical regions.

**Figure A1.**
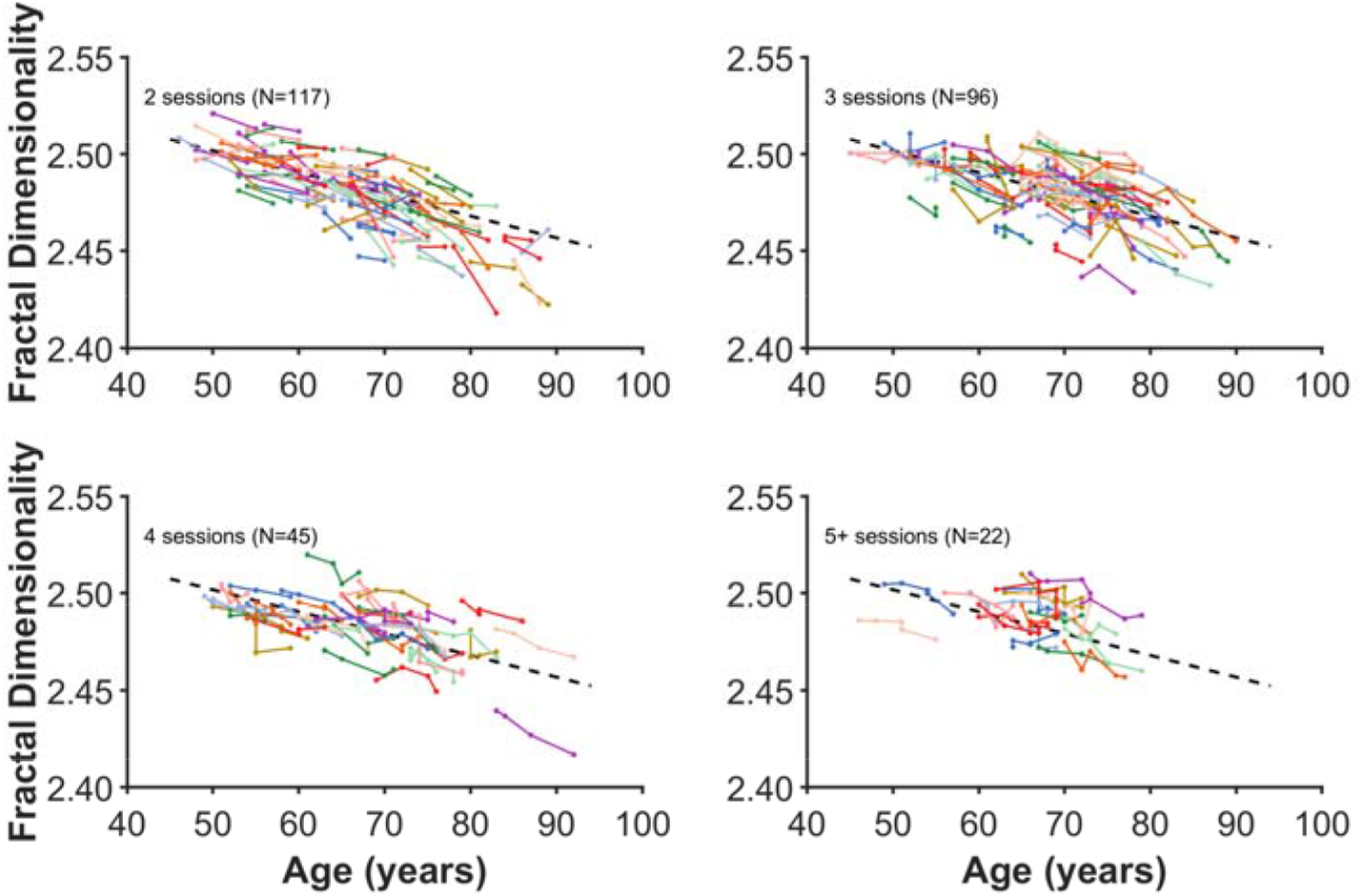
Age changes in global cortical fractal dimensionality. Each coloured line represents an individual participant, with each dot corresponding to an MRI session. The plot has been divided into panels for each set of timepoints to improve readability. Dashed line present in all panels shows the decrease in global fractal dimensionality based on all data, estimated using the linear mixed effects model. Compare with Figure 3.

